# Assessment of AI-based Protein Structure Prediction for the NLRP3 Target

**DOI:** 10.1101/2021.11.05.467381

**Authors:** Jian Yin, Junkun Lei, Jialin Yu, Weiren Cui, Yifan Zhou, Hua Feng, Jason Deng, Wenji Su, Letian Kuai

## Abstract

The recent success of AlphaFold and RoseTTAFold has demonstrated the values of AI methods in predicting highly accurate protein structures. Despite the advances, their roles in the context of small-molecule drug discovery need to be thoroughly explored. In this study, we evaluated whether the AI-based models can lead to reliable three-dimensional structures of protein-ligand complexes. The structure we predicted was NLRP3, a challenging protein target in terms of obtaining the three-dimensional model both experimentally and computationally. The conformation of the binding pockets generated by the AI models were carefully characterized and compared with experimental structures. Further molecular docking results indicated that AI-predicted protein structures combined with molecular dynamics simulations can be useful in small-molecule drug discovery.

## Introduction

High-resolution X-ray crystallographic or NMR structure of proteins and protein-ligand complexes are critical for understanding the interactions between small-molecule drug candidates and their targets, and therefore are pivotal in the drug discovery process.^[1]^ However, due to the limitations of experimental techniques, there are still a large fraction of protein structures that cannot be solved. Studies on novel drug targets are often hampered by a lack of protein structures with sufficient resolution or completeness.

The AI methods for protein structure predictions, including AlphaFold (AF) ^[2]^ and RoseTTAFold (RF) ^[3]^, have attracted tremendous interest for their ability to accurately predict protein structures, without relying on structural templates with high sequence similarity. Albeit the success in recent evaluations of single protein structure predictions, their applications in small-molecule drug discovery have not been systematically investigated yet. For instance, protein targets may undergo induced-fit conformational changes due to the binding of small molecules^[4]^. Predictions might become even more challenging when the protein comprises multiple domains or subdomains. ^[5]^

In the current study, we explored whether the AI predicted protein structures can be employed to predict the binding modes of known NLRP3 inhibitors. The target NLRP3 (NOD-like receptor family, pyrin domain-containing protein 3) ^[6]^ is a protein linked to many chronic inflammatory human diseases, such as atherosclerosis, Alzheimer’s disease, and nonalcoholic steatohepatitis. It is part of the NLRP3 inflammasome that, when activated, triggers the release of proinflammatory cytokines IL-1β and IL-18, and thus lead to an inflammatory form of cell death^[7]^. Despite the considerable interest in this target, a lack of structural information had hindered the development of small-molecule inhibitors. We therefore carefully evaluated the quality of the AI predicted models, and assessed the discrepancy between the binding pockets formed in AI models and the experimental structures. Furthermore, we demonstrated that the AI models combined with molecular dynamics (MD) simulations are valuable for exploring the mechanisms of small-molecule drug candidates and facilitating lead optimization process.

## Results

### Evaluation of the AI-predicted models of NLRP3

The human NLRP3 protein contains subunits PYD (pyrin domain; Residues 3-95), FISNA (Fish-specific NACHT associated domain; 96-218), LRR (leucine-rich repeat domain; 652-1036) and a central NACHT domain comprising NBD (nucleotide-binding domain; 219-372), HD1 (helical domain 1; 373-434), WHD (winged helix domain; 435-541) and HD2 (helical domain 2; 542-651) (Fig. 1). For a long time, the only available experimental structure containing the central subdomains of this protein was a cryo-electron microscopy (cryo-EM) model bound to NEK7, a mitotic kinase mediating NLRP3 activation (PDB code: 6NPY; Fig. 1a). A substantial portion of the residues (31.8%) in the HD2 region near the small molecule binding site in this structure was missing. In addition, the conformation of NLRP3 complexed with the NEK7 substrate may differ from that when bound to a small-molecule inhibitor. A cryo-EM structure of NLRP3 decamer bound to drug MCC950 (PDB code: 7PZC) was newly released on January 26, 2022. ^[8]^ Besides, a crystal structure with a MCC950-like inhibitor (PDB code: 7ALV) was released on October 20, 2021 ^[9]^. But note that those two structures had not yet been deposited to PDB when we made structural predictions in the current study.

**Figure 1.**
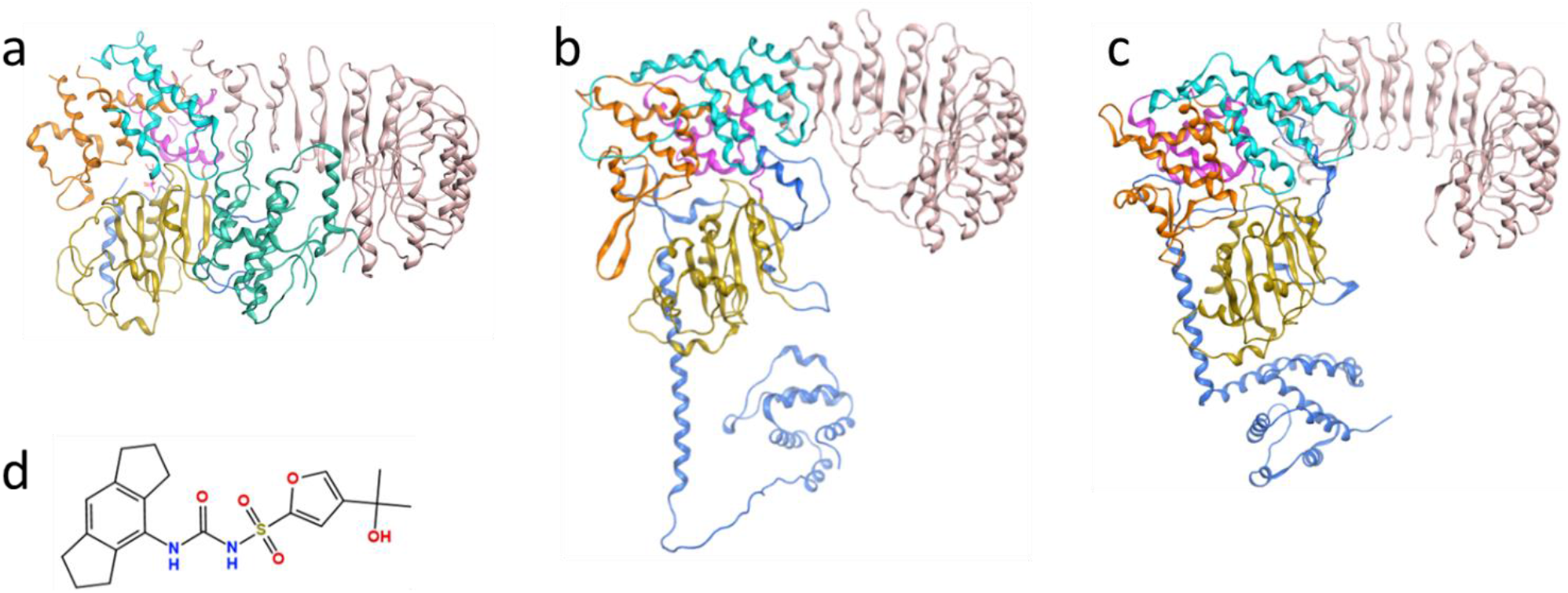
(a) Previously reported cryo-EM structures of NLRP3 complexed with NEK7 kinase (PDB code: 6NPY); the structures of NLRP3 predicted by (b) AlphaFold and (c) RoseTTAFold. Gold: NBD; Purple: HD1; Green: NEK7; Orange: WHD; Cyan: HD2; Pink: LRR; Blue: others; (d) the 2D structure of MCC950, which was reported as a selective NLRP3 inhibitor.

Given the difficulties in obtaining the structure of the multi-subdomain NLRP3 protein using experimental and traditional theoretical methods, we chose this protein to test the accuracy of the AF and RF models for predicting protein conformations, and whether correct protein-small molecule binding modes can be generated based on these structures. Five models were generated by two locally installed programs each. The pLDDT (predicted Local Distance Difference Test) scores^[2]^ were calculated for every global structure and subdomain to quantify the confidence in structure prediction, which were interpreted as follows: >90: high accuracy; 70–90: moderate to good; 50-70: low confidence; <50: disordered or unstructured. It was observed that the AF and RF models showed comparable overall confidence level, with an average value of 79.9 and a standard deviation of 0.9 for AlphaFold, compared with 78.3 ±1.2 for RoseTTAFold. It is also noteworthy that all pLDDT scores of the central subdomains are above 70, indicating that the protein was reliably predicted at the individual domain level.

For multi-domain or -subdomain protein targets such as NLRP3, the predicted templated modelling (pTM) score ^[2]^ can be used to measure the accuracy of domain packing. The AF package allows us to obtain the pTM scores from the fine-tuned pTM models (Table 1). The numbers around 0.7 indicated moderate risk associated with incorrect domain assembly and inter-domain contacts, which should be of particular concern because the binding site of the known inhibitor MCC950 is located at the interfaces of several central subdomains.

**Table 1.** Predicted Local Distance Difference Test (pLDDT) and predicted Templated Modelling (pTM) scores for the AI-predicted structures of the NLRP3 protein.

| Domain | Model1 | Model2 | Model3 | Model4 | Model5 |
| --- | --- | --- | --- | --- | --- |
| NBD | 83.3 | 85.3 | 86.8 | 83.5 | 85.7 |
| HD1 | 82.8 | 84.9 | 85.4 | 82.1 | 84.3 |
| WHD | 70.2 | 71.5 | 74.2 | 71.0 | 71.2 |
| HD2 | 76.8 | 78.6 | 79.4 | 77.2 | 79.5 |
| LRR | 84.6 | 85.0 | 86.4 | 85.3 | 84.7 |
| Full-length | 79.1<br>(0.58) | 80.3<br>(0.67) | 81.4<br>(0.71) | 79.1<br>(0.70) | 79.8<br>(0.71) |
| NBD | 83.4 | 81.6 | 82.5 | 80.7 | 79.5 |
| HD1 | 83.5 | 82.3 | 81.3 | 80.7 | 80.3 |
| WHD | 80.0 | 78.4 | 76.4 | 78.7 | 77.3 |
| HD2 | 75.2 | 73.1 | 70.2 | 73.0 | 71.1 |
| LRR | 82.3 | 82.9 | 83.1 | 81.9 | 81.8 |
| Full-length | 79.7 | 79.4 | 77.9 | 77.6 | 76.8 |

\*pLDDT scores are reported on a 0 -100 scale. Values higher than 90: high accuracy; 70–90: moderate to good; 50–70: low confidence; <50: disordered or unstructured. pTM scores (numbers in the parentheses)
are on a 0-1 scale. Numbers higher than 0.7 indicate high accuracy.

Next, we computed the RMSD values between the predicted models and one of the monomers in the experimental NLRP3 decamer (PDB code: 7PZC). As shown in Table 2, the RF model performed better on WHD, while the prediction of the AF model at the HD2 region was closer to the experimental structure. In addition, the RMSD values of the LRR subdomain structures were significantly higher than those of other subdomains, using both AF and RF methods, resulting in an overall RMSD above 10 Å for all models. To focus on subdomains and residues that play critical roles in small-molecule binding, we then performed a detailed analysis of the differences between the AI models and the experimental structures at the MCC950 binding sites. The results are described in the following section.

**Table 2.** RMSD values between the AI-predicted NLRP3 structures and one of the monomers in 7PZC (Unit: Å).

| Domain | Model1 | Model2 | Model3 | Model4 | Model5 | # of aligned atoms |
| --- | --- | --- | --- | --- | --- | --- |

| AlphaFold |  |  |  |  |  |  |
| --- | --- | --- | --- | --- | --- | --- |
| NBD | 2.5 (1.7) | 2.6 (1.7) | 2.5 (1.7) | 2.6 (1.8) | 2.6 (1.8) | 1227 (154) |
| HD1 | 2.4 (1.5) | 2.4 (1.5) | 2.7 (2.0) | 2.2 (1.3) | 2.4 (1.5) | 505 (62) |
| WHD | 5.7 (4.9) | 5.9 (5.2) | 5.8 (5.1) | 5.7 (4.9) | 5.8 (5.1) | 873 (107) |
| HD2 | 4.4 (3.6) | 4.4 (3.5) | 4.1 (3.1) | 4.0 (3.1) | 4.2 (3.4) | 921 (110) |
| LRR | 11.1 (10.4) | 11.8 (11.1) | 9.5 (8.9) | 4.4 (3.8) | 11.5 (10.8) | 2945 (385) |
| Full-length | 25.2 (24.8) | 26.6 (26.1) | 18.8 (18.3) | 10.6 (10.2) | 17.4 (17.0) | 8048 (1010) |
| RoseTTAFold |  |  |  |  |  |  |
| NBD | 2.9 (1.9) | 2.8 (1.9) | 2.8 (1.9) | 2.8 (1.9) | 2.8 (1.9) | *Same as*<br>*AlphaFold* |
| HD1 | 2.3 (1.3) | 2.4 (1.4) | 2.4 (1.3) | 2.4 (1.3) | 2.3 (1.3) |  |
| WHD | 3.5 (2.1) | 3.5 (2.1) | 3.5 (2.1) | 3.4 (2.0) | 3.4 (2.1) |  |
| HD2 | 8.0 (7.5) | 5.3 (4.3) | 5.2 (4.2) | 7.2 (6.6) | 7.1 (6.5) |  |
| LRR | 10.9 (10.6) | 10.9 (10.6) | 11.0 (10.7) | 10.9 (10.7) | 11.1 (10.7) |  |
| Full-length | 13.0 (12.5) | 14.8 (14.4) | 23.8 (23.3) | 21.9 (21.3) | 21.8 (21.5) |  |

\*Numbers before and in the parentheses are the RMSD values calculated using coordinates of all heavy atoms and $\alpha$ -carbons, respectively. Note that the RMSD of the subunits were computed by superposing each set of predicted and experimental subdomains separately.

### Characterization of the Binding pocket of compound MCC950

MCC950 has long been identified as a selective NLRP3 inhibitor that blocks NLRP3 inflammasome activation ^[10, 11]^ (Fig. 1d), but the mechanism was unclear until very recently. The newly reported experimental structure showed that MCC950 was bound to a cleft formed by subdomains NBD, HD1, WHD, HD2 and LRR ^[8]^, which was separated from the ADP binding site by the Walker A motif (GAAGIGKTIL)^[12]^. The residues adjacent to MCC950 include F575, R578 and E629 in the HD2 subdomain, A228 in the Walker A motif of NBD, as well as I411 and Y443 in HD1 (Fig. 2 and 3a). Therefore, we measured the distances among the α-carbon atoms of these key residues in each AI model, as well as the distances among their side chains, to assess the possibility of MCC950 binding pocket formation. In particular, the carbamide and sulfonyl groups of MCC950 engaged hydrogen bonds with residues A228 and R578, residing in different subdomains across the binding pocket (Fig. 4a). As a result, the distance between A228 and R578 residues was considered as one of the key indicators for the formation of a suitable binding site.

**Figure 2.**
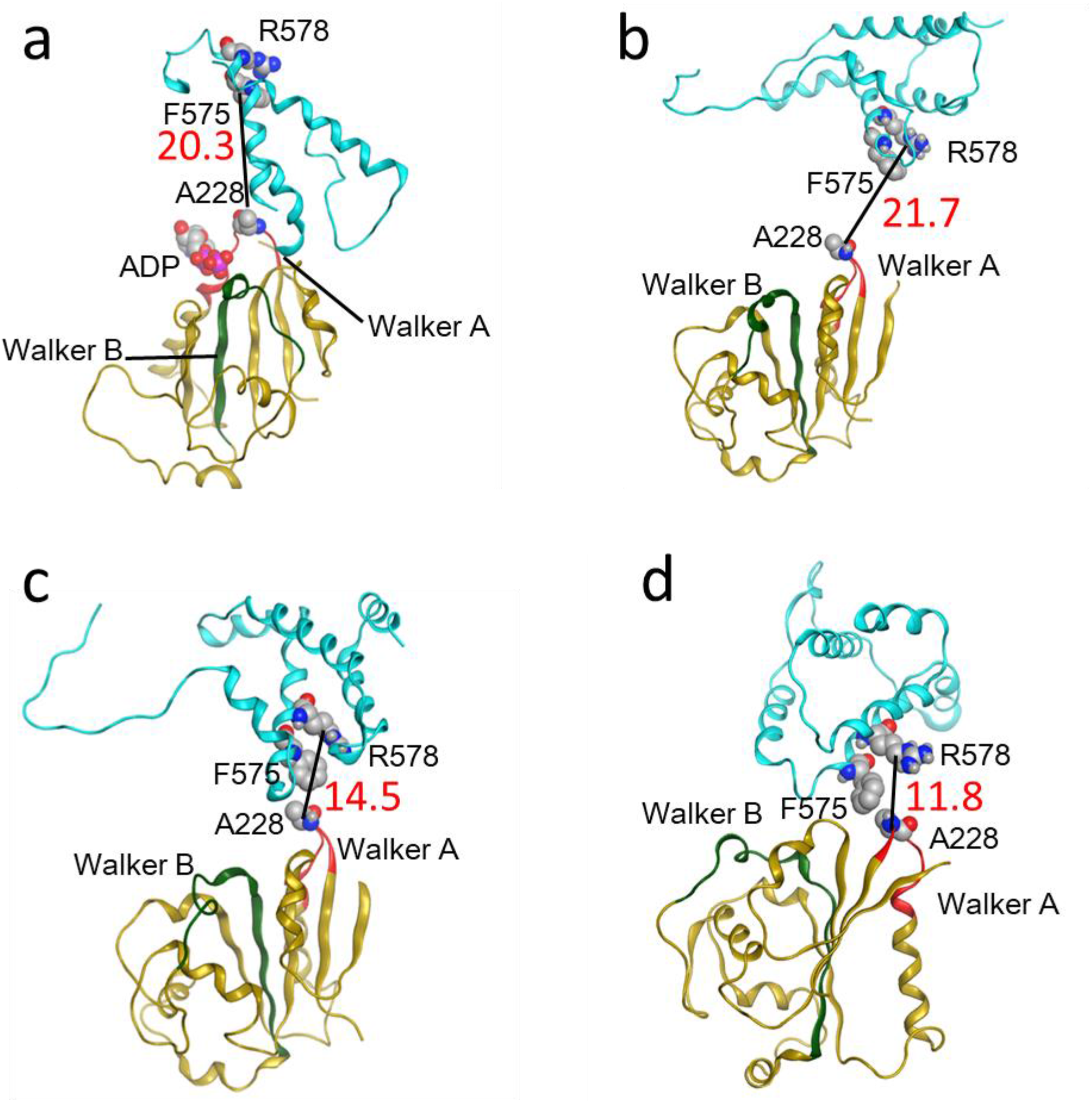
Relative positions between two key residues R578 and A228 of the MCC950 binding pocket in (a) the NLRP3-NEK7 complex structure (PDB code: 6NPY), (b) model 1 generated by AlphaFold, (c) model 4 by AlphaFold; and (d) model 1 by RoseTTAFold. The distance between the α-carbon atoms of R578 and A228 was 20.3 Å, 21.7 Å, 14.5 Å and 11.8 Å in (a), (b), (c) and (d), respectively. Red: the Walker A motif (GAAGIGKTIL); Green: the Walker B motif previously suggested as the binding region of MCC950 (RILFMDGFDELQGAFDEHI)[^12^]; Gold: NBD; Cyan: HD2. Other domains omitted for clarity.

**Figure 3.**
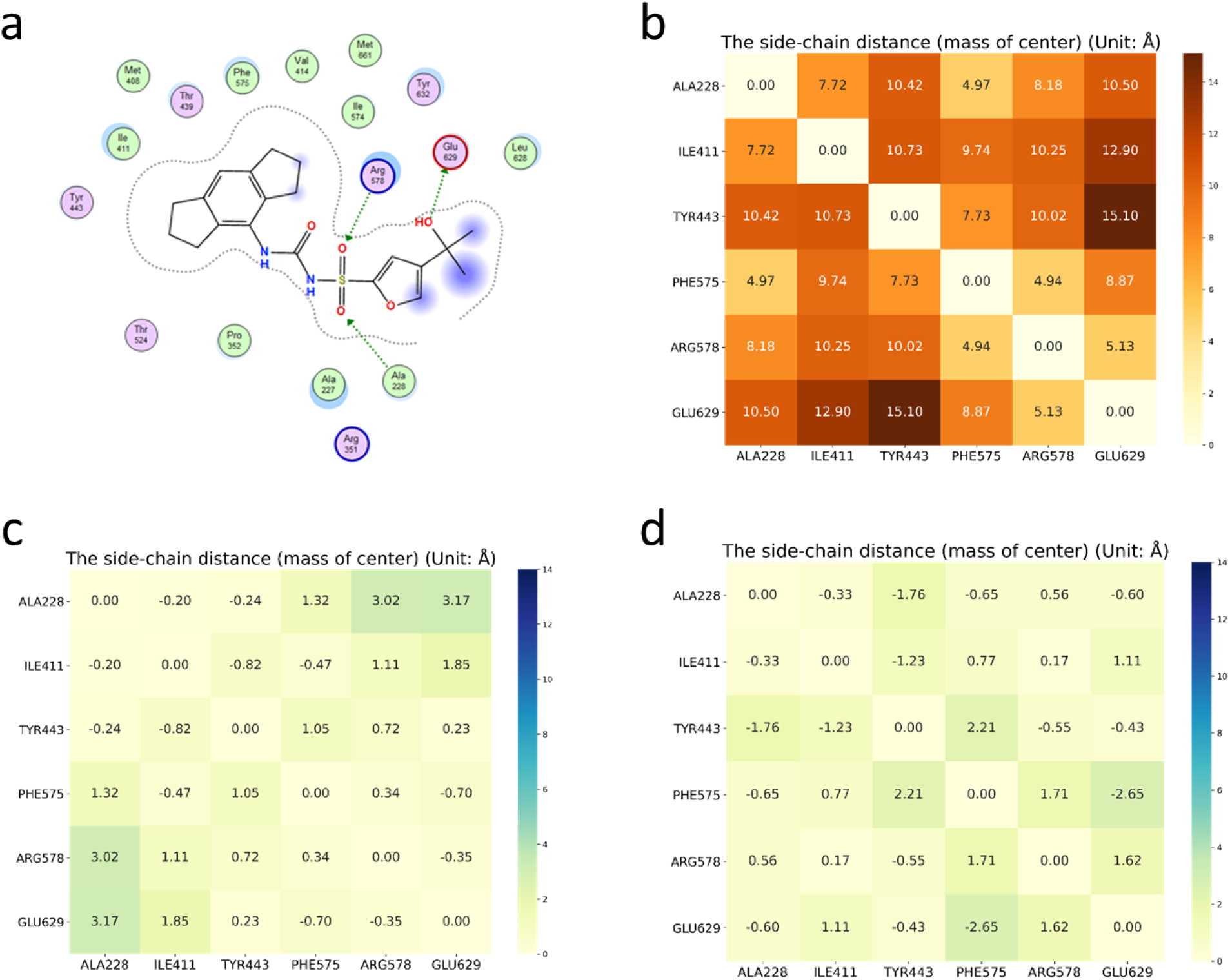
(a) Major interactions between MCC950 and NLRP3 in 7PZC; (b) side-chain distances between pairs of key residues at the binding site of MCC950 in 7PZC; side-chain distances between pairs of key residues in (c) AF model 4 and (d) RF model 1 relative to those in 7PZC.

**Figure 4.**
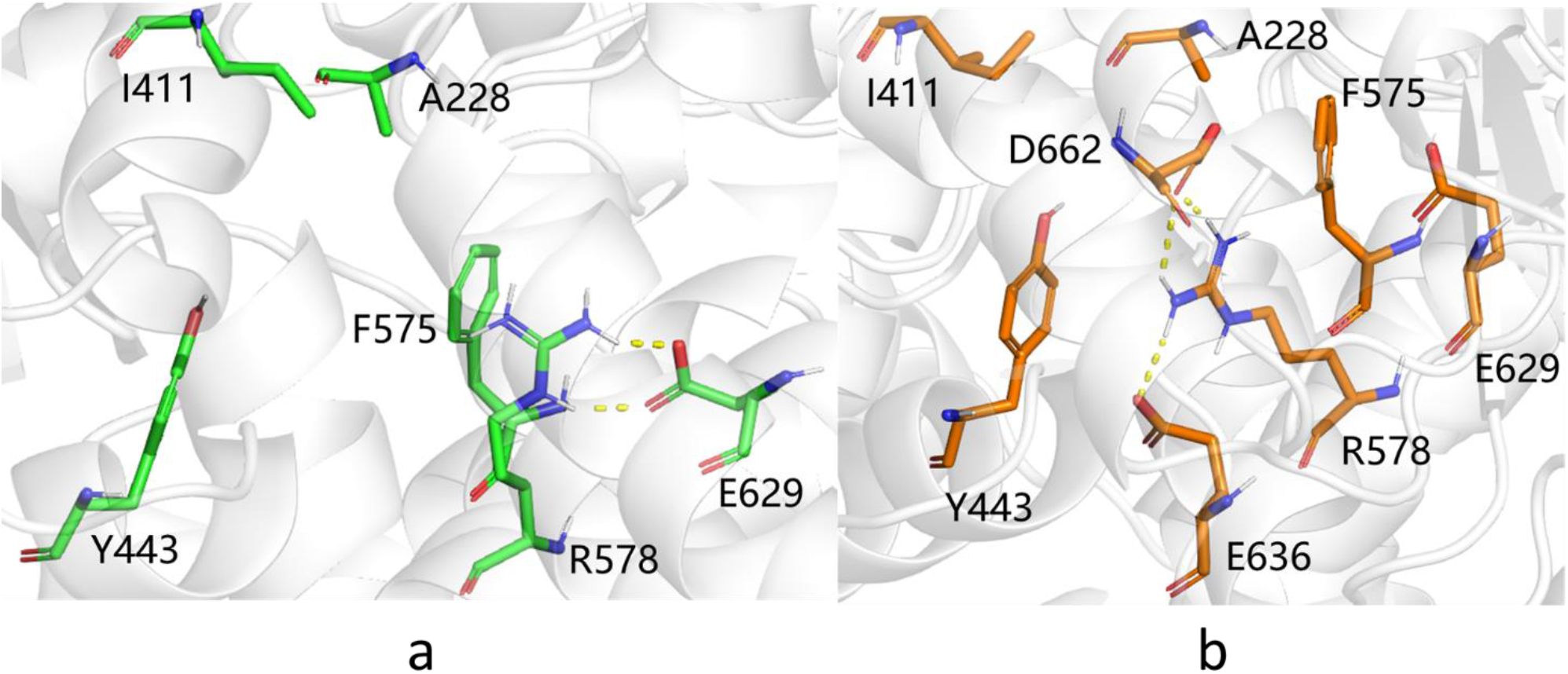
Salt bridges in the MCC950 binding pocket of (a) AF model 4 and (b) RF model 1.

The results showed that R578 and its adjacent residue F575 in the experimental structure of the NLRP3-NEK7 complex (PDB code: 6NPY) were located in the solventexposing region, relatively distant from the ADP-binding site (Fig. 2a). The distance between R578 and A228 was measured as 20.3 Å. In the AF models, the R578 and A228 residues were placed facing each other, yet the distance between them was around 20 Å in each AF structure except for model 4 with a value of 14.5 Å, which indicated that the HD2 and NBD subdomains were not sufficiently packed together (Fig. 2b and 2c), considering the corresponding distance of 12.7 Å in the experimental structure 7PZC. Note that the RMSD between AF model 4 and the experimental structure was also computed as the smallest value among all AI models (Table 2). In contrast, the HD2 and NBD subdomains in the RF models were more densely packed, and the R578 and A228 residues at the interfaces seemed close enough (~12 Å) to interact with the small-molecule inhibitor simultaneously (Fig. 2d).

In addition to the hydrogen bonds formed between the amide group of MCC950 and the protein, the tricyclic head group of the small molecule also participated in hydrophobic interactions with neighboring residues involving I411, Y443, and F575. Besides, the terminal carboxylate group engaged H-bonds with G629 (Fig. 4a). We therefore compared the pairwise distances of related residues for all AF and RF models with those in 7PZC, as a finer measure of binding pocket structural integrity. It was observed that, on average, AF model 4 had the smallest discrepancy from the experimental results among all AI models (Fig. 3). But still, the green areas shown in Fig. 3c suggested a loosely packed core. In particular, similar salt bridge between R578 and E629 was observed in the AF models and 7PZC (Fig. 4a and 6a), which further stabilized the complexation between the small molecule and the protein, whereas in the RF models, R578 instead formed salt bridges with two other negatively charged residues E636 and D662 (Figure 4b).

To further assess the use of AI models in small molecule drug discovery, we performed molecular docking of MCC950 for each model of AF and RF. However, even the top-ranked ones produced only partially correct binding modes. As a result, molecular dynamics (MD) simulations were carried out to refine poses by taking account of the induced fit and protein dynamic effects. To keep the Walker A moiety from shifting, ADP was docked back into its binding site of the AI models, and then simulated by MD for 200 ns. It turned out the RF model 1 structure could hardly accommodate ADP at the corresponding site. As a result, we only performed a 200 ns MD simulation of AF model 4 bound to ADP (Fig. 5a). Another difficulty was that part of the FISNA domain missing from the experimental structure in AF Model 4 occupied the entrance of the binding pocket. Therefore, ligand MCC950 was docked to the protein-ADP conformation from the last frame of the simulation with the PYD and FISNA subdomains truncated, and then simulated for another 200 ns (Fig. 5b).

**Figure 5.**
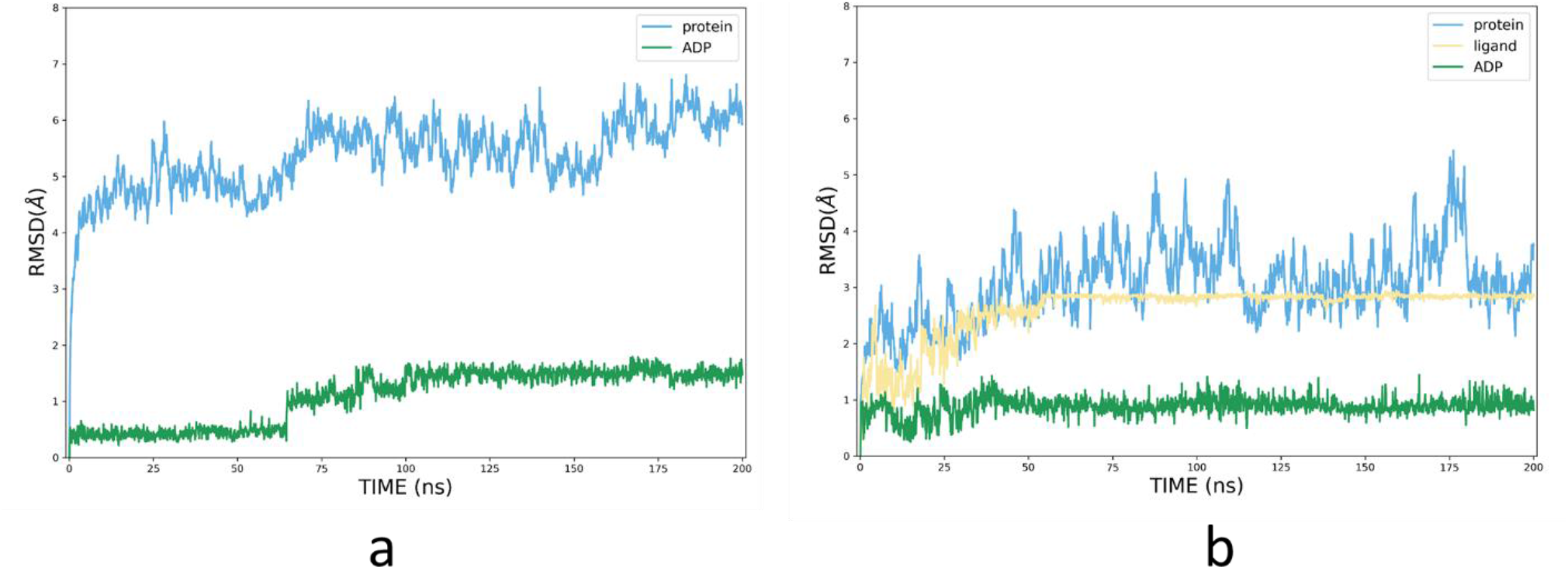
RMSD values plotted relative to the first frame (in Å) of the production phase based on the MD trajectories of (a) AF model 4 bound to ADP and (b) AF model 4 bound to ADP and MCC950. Blue line: *C_α_* RMSDs of the protein; green and yellow lines: heavy atom RMSDs of ADP and the ligand, respectively.

Compared with the experimental structure, ligand MCC950 engaged similar hydrogen-bonding interactions with R578, A228 and G629 (Fig. 6). The distance between the α-carbon atoms of R578 and A228 was reduced to 14.1 Å from the initial 14.5 Å, which became closer to the measured value of 12.7 Å in 7PZC. It was challenging to fully optimize the assembly of subdomains through MD simulations because conformational changes involving the main chain typically require a long time to converge.

**Figure 6.**
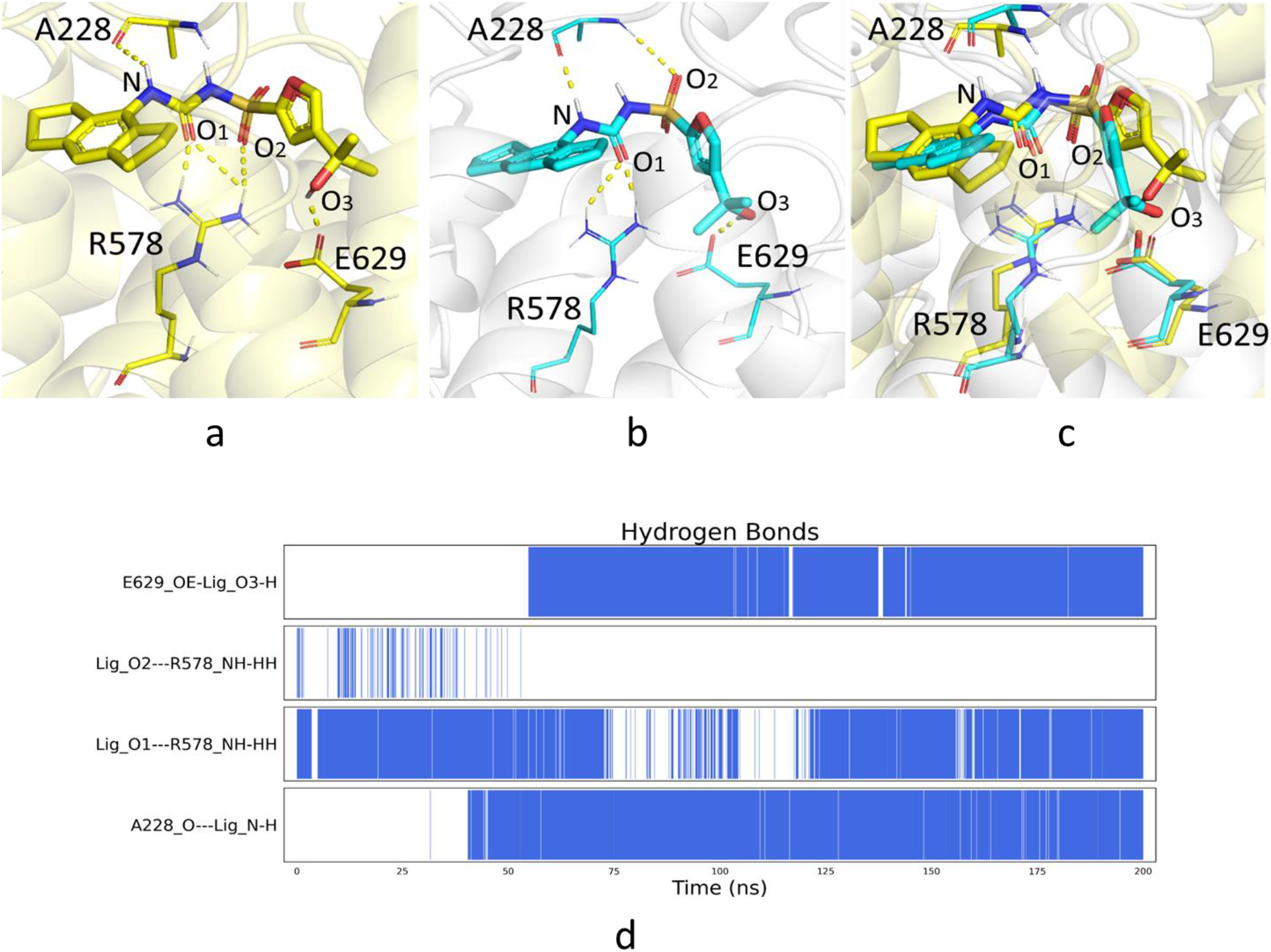
The binding modes of MCC950 in (a) 7PZC and (b) AF model4 in complex with ADP and MCC950 from the last frame of the 200 ns MD simulations; (c) structural overlay of (a) and (b); (d) Hydrogen bond analysis of the MD simulations.

## Discussion

Despite the use of protein structure templates during both training and predicting procedures, AlphaFold and RoseTTAFold are intrinsically more advanced than the conventional template-based methods such as homology modelling and threading. With the neural network architecture, the AI methods are able to learn deeply from the coevolution of protein sequences, and detect the underlying correlations between pairs of amino acids through the construction of multiple sequence alignment (MSA) and pair representation, instead of merely patching the templates together. Therefore, the AI methods can accurately predict protein structures using templates with sequence identity and structural coverage lower than 30%. Indeed, this current study demonstrated that the structures predicted by the AI methods did not heavily rely on the existing experimental structure, and could be a good complementary method for experimental techniques including X-ray crystallography, NMR and cryo-EM technique.

Accurately predicting the protein structure in the present work is a particularly challenging case because the NLRP3 protein consists of multiple subdomains, and the small-molecule binding pocket is located in the interdomain region. It was shown that both AlphaFold and RoseTTAFold can predict confidently at the subdomain level. Nevertheless, the subdomains were more compact in RoseTTAFold models in the present study, whereas AlphaFold successfully reproduced a key salt bridge interaction observed in the experimental structure. Our study suggested that one should be careful when using AI protein structure prediction tools for drug targets comprising of two or more structural domains, especially when the drug binding site is formed at the interfaces of subunits. MD simulations can be applied to finetune the details of models in complex systems. In practice, AI predicted models that can be correlated with the results of affinity selection experiment may be regarded as more trustworthy. The effort of validating AI generated protein structures based on affinity selection data are currently ongoing.

### Simulation Details

The docking poses were obtained using MOE ^[13]^. All MD simulations were performed using AMBER20^[14]^ program. Each system was solvated with TIP3P^[15]^ water molecules in a rectangular box with a 10 Å buffer distance. Partial atomic charges of small molecules were derived using AM1-BCC^[16]^. Bonded and Lennard-Jones parameters were obtained from GAFF2^[17]^. The Amber force field ff19SB^[18]^ was used for the protein. Counterions were only added to neutralize the total charge of each system.

The whole system was first minimized using a steepest descent algorithm, with the maximal number of cycles set as 10 000. Subsequently, the system was heated from 297 K to 300 K over a period of 50 ps, and this was followed by a 100 ps NVT and a 200 ps NPT equilibrations with all heavy atoms restrained during this process.

The production phase was simulated in an NPT ensemble with the Berendsen barostat^[19]^ and Langevin^[20]^ thermostat at 300 K without restraints. The cutoff distance of Lennard-Jones interactions was set to 10 Å and a long-range correction was applied to approximate the interactions beyond the cutoff distance. Electrostatic interactions were treated with the particle mesh Ewald (PME) scheme^[21]^. The simulation time of the production run was 200 ns for both the ADP-bound protein and the subsequent MCC950 docked complex.

## Supplementary Materials

Raw models generated by AlphaFold and RoseTTAFold are provided. When querying templates for the protein target, we used the PDB downloaded on May 14, 2020 and the PDB70 cluster database downloaded on May 13, 2020, both before the release dates of the experimental structures 7PZC and 7ALV.

